# High-content high-resolution microscopy and deep learning assisted analysis reveals host and bacterial heterogeneity during *Shigella* infection

**DOI:** 10.1101/2024.03.06.583762

**Authors:** Ana T. López-Jiménez, Dominik Brokatzky, Kamla Pillay, Tyrese Williams, Gizem Özbaykal Güler, Serge Mostowy

## Abstract

*Shigella flexneri* is a Gram-negative bacterial pathogen and causative agent of bacillary dysentery. *S. flexneri* is closely related to *Escherichia coli* but harbors a virulence plasmid that encodes a Type III Secretion System (T3SS) required for host cell invasion. Widely recognized as a paradigm for research in cellular microbiology, *S. flexneri* has emerged as important to study mechanisms of cell-autonomous immunity, including septin cage entrapment. Here we use high-content high-resolution microscopy to monitor the dynamic and heterogeneous *S. flexneri* infection process by assessing multiple host and bacterial parameters (DNA replication, protein translation, T3SS activity). In the case of infected host cells, we report a reduction in DNA and protein synthesis together with morphological changes that suggest *S. flexneri* can induce cell-cycle arrest. We developed an artificial intelligence image analysis approach using Convolutional Neural Networks to reliably quantify, in an automated and unbiased manner, the recruitment of SEPT7 to intracellular bacteria. We discover that heterogeneous SEPT7 assemblies are recruited to bacteria with increased T3SS activation. Our automated microscopy workflow is useful to illuminate diverse host and bacterial interactions at the single-cell and population level, and to fully characterise the intracellular microenvironment controlling the *S. flexneri* infection process.

## Introduction

*Shigella flexneri* is a Gram-negative bacterium and the etiologic agent of bacillary dysentery, an intestinal infectious disease that kills more than 150 thousand people worldwide per year, disproportionally affecting children under 5 years old from low- and middle-income countries (Global Burden of Disease Collaborative Network., 2020). After the ingestion of a small number of bacteria (10-100), *S. flexneri* rapidly colonises the human intestinal epithelium. For this, *S. flexneri* uses a Type 3 Secretion System (T3SS) that injects bacterial effectors into the host cell to modulate key aspects of the infection process and to evade innate immune defences by the host (Schnupf & Sansonetti, 2019). Following uptake by host cells, *S. flexneri* breaks its vacuolar compartment to access the cytosol (Ray et al., 2010). Here, the virulence factor IcsA enables *S. flexneri* to hijack host actin and polymerise comet tails that render bacteria motile to disseminate to neighbouring cells and spread in the tissue (Bernardini et al., 1989; Egile et al., 1999).

Septins are highly-conserved GTP-binding proteins that form heteromeric complexes (hexamers, octamers) that assemble into non-polar filaments, bundles and rings (Martins et al., 2023; Soroor et al., 2021; Szuba et al., 2021; Woods & Gladfelter, 2021). Septins are involved in many physiological cellular processes (eg. cytokinesis, cilia formation), and play key roles during infection (Mostowy & Cossart, 2012; Robertin & Mostowy, 2020; Spiliotis & Nakos, 2021). Septins form collar-like structures at the plasma membrane during bacterial invasion (Boddy et al., 2018; Mostowy et al., 2009; Robertin et al., 2023), and ring-like structures surrounding membrane protrusions in cell-to-cell bacterial spread (Mostowy et al., 2010). In the cytosol septins are recruited to actin-polymerising bacteria, including *S. flexneri* and *Mycobacterium marinum*, as cage-like structures (Mostowy et al., 2010). In the case of *S. flexneri*, septins are recruited to µm-scale curvature at bacterial poles and the division site, which are enriched in the negatively charged phospholipid cardiolipin (Krokowski et al., 2018). Recent work has shown that IcsA promotes the recruitment of septins to *S. flexneri*, and that lipopolysaccharide (LPS) inhibits *S. flexneri*-septin cage entrapment (Lobato-Márquez et al., 2021; Mostowy et al., 2010). Septin cages prevent bacterial actin tail motility and therefore are considered as an antibacterial mechanism of host defence (Krokowski et al., 2018; Mostowy et al., 2010).

Intracellular infection is usually portrayed as a uniform, synchronised and highly orchestrated process. However, infection assays performed in controlled laboratory environments with isogenic populations display a high level of complexity (Avraham et al., 2015; Toniolo et al., 2021). It is particularly unknown how different subsets of bacteria (eg. motile vs. non motile bacteria) can colonise distinct intracellular microenvironments and become targeted by multiple cell-autonomous immunity mechanisms (eg. septin cages, autophagy). It has been proposed that this diversity of intracellular niches impacts bacterial physiology (Day et al., 2023; Gutierrez & Enninga, 2022; Santucci et al., 2021). For example, it may affect individual bacterial ability to acquire nutrients, to be exposed to varying antibiotic concentrations, or to elicit different responses and signalling pathways within the host cell. While the concept of phenotypic heterogeneity in prokaryotic clonal populations is well established (Ackermann, 2015), the concept is emerging also in eukaryotic cells where non genetic differences may for instance explain individual cell sensitivity and responses by cell-autonomous immunity (McDonald & Dedhar, 2024). In addition, previous high-content studies coupled with mathematical modelling have shown that host cell context (including local cell density, lipid composition at the plasma membrane, or previous infection status) affects their susceptibility to viral and bacterial infection (Snijder et al., 2009; Voznica et al., 2018). It is therefore paramount to study host and bacterial cells at the single-cell level, to assess the individual responses that collectively contribute to the infection process.

Here, to dissect the complexity of *S. flexneri* infection in epithelial cells, we use high-content high-resolution microscopy coupled to automated image analysis. High-content microscopy enables the automated acquisition of thousands of microscopy images and therefore is useful to capture the heterogeneity of individual host and bacterial cells during the infection process for quantitative analysis (Aylan et al., 2023; Brodin & Christophe, 2011; Deboosere et al., 2021; Dramé et al., 2023; Fisch et al., 2019a; Lensen et al., 2023; Pylkkö et al., 2021). We analyse multiple parameters, such as morphological host cell features, the *de novo* synthesis of DNA and proteins in both host and bacterial cells, as well as activation of the bacterial T3SS. To investigate the antimicrobial potential of septin-mediated immunity, we develop a deep-learning based approach to automatically identify septin-*S. flexneri* interactions. Together, we highlight the power of high-content microscopy and artificial intelligence to reveal insights on both host and bacterial cells, and illuminate fundamental biology underlying the *S. flexneri* infection process.

## Results

### Capturing the heterogeneity of *S. flexneri* infection using high-content microscopy

To understand the complexity and reveal the heterogeneity of *S. flexneri* infection in human epithelial cells, we dissected the infection process using high-content high-resolution microscopy and HeLa cell infection model (**Figure 1A, B, Figure 1-figure supplement 1A**). We observed a mean of 11.5% of infected cells measured by fluorescence at 3h 40 min post infection (hpi) at MOI 100:1 (**Figure 1C**). These infected cells presented different burdens of bacteria following an exponential decay distribution (**Figure 1D**), reflective of the formation of infection foci (Ortega et al., 2019). With growing evidence of the cytosol being a non-homogeneous compartment containing concentration gradients, biomolecular condensates, protein complexes and concentrates (van Tartwijk & Kaminski, 2022), we tested whether *S. flexneri* may be exposed to different intracellular microenvironments. For this, we measured the distance of bacteria to the centroid of cells (**Figure 1E**) and observed a median distance of 8.74 µm. The median distance to the centroid of the cell significantly decreased with increasing bacterial burden (**Figure 1E**), suggesting that the microenvironment of bacteria is diverse and changes as bacteria replicate intracellularly and populate the central parts of host cytosol after uptake at the host cell periphery. To dissect the heterogeneity in host cells, we quantified an increase of 36.9% and 22.3% in the cellular and nuclear area of infected cells, respectively (**Figure 1G, I**), which was dependent on the infection burden (**Figure 1H, J**). In addition, the total amount of Hoechst fluorescence in the cell nucleus increased 29.4% in infected cells (**Figure 1-figure supplement 1B**), indicating either an increase in nuclear volume or altered chromatin condensation. To test if this effect was due to morphological rearrangements during infection, we treated cells with Latrunculin B to inhibit actin polymerisation. While cells treated with Latrunculin B decreased their cellular area (but not their nuclear area) (**Figure 1-figure supplement 1C, D**), the drug did not abrogate the difference in size between infected and uninfected cells. Taken together, these results highlight profound morphological rearrangements during infection that are dependent on the infection burden.

**Figure 1.**
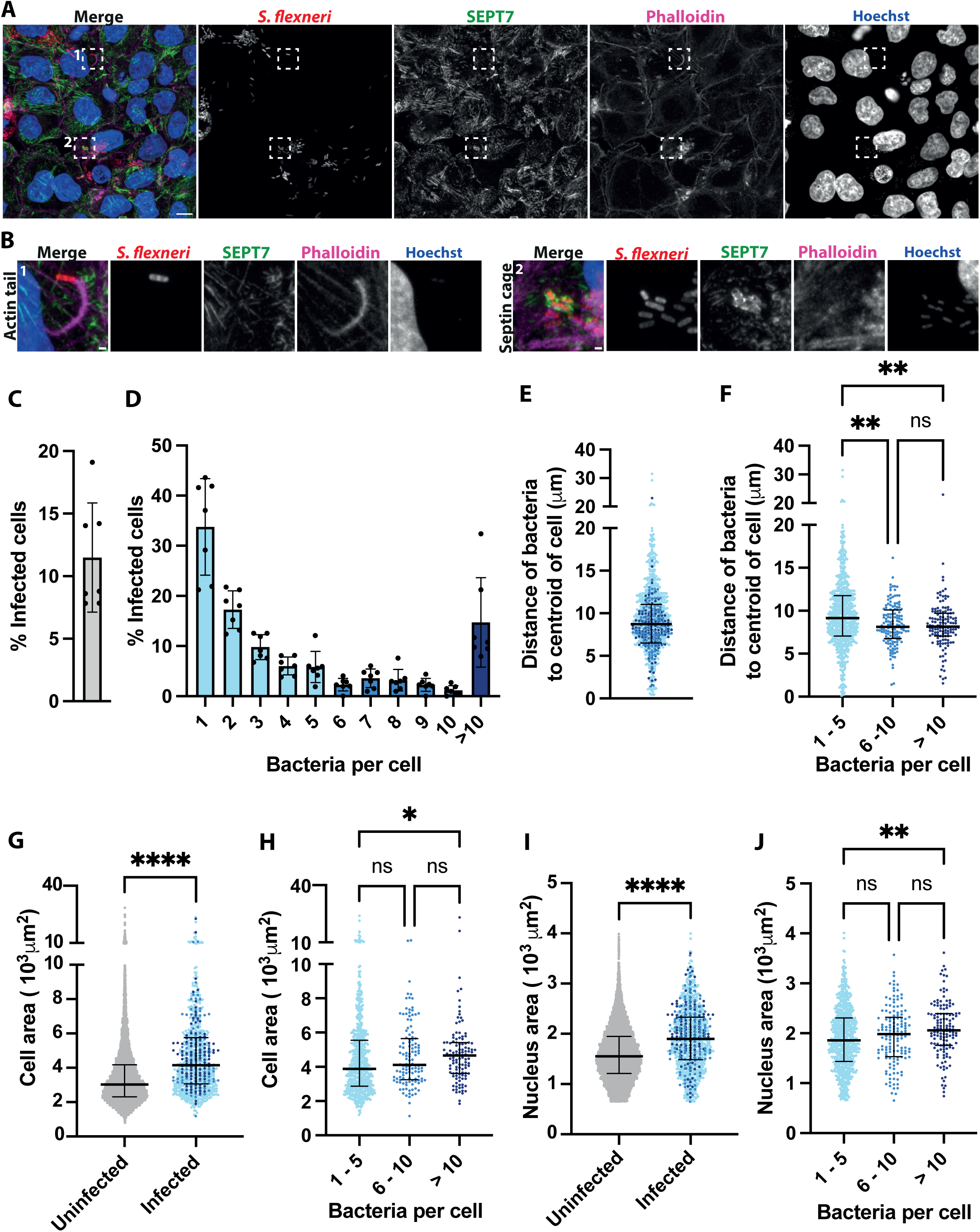
High-content imaging captures the heterogeneity of *S. flexneri* infection. (A) Representative high-resolution microscopy image of HeLa cells infected with *S. flexneri* expressing mCherry acquired with the high-content microscope ZEISS CellDiscoverer7. Scale bar, 10 µm. (B) Enlarged view of insets in A, highlighting a *S. flexneri* bacteria polymerising an actin tail (1) and entrapped in a septin cage (2), scale bar, 1 µm. (C) Percentage of infected cells at 3 h.p.i. of 7 independent experiments. Graph represents mean = 11.49% ± SD = 4.36%. (D) Percentage of infected cells harbouring different bacterial doses, graph represents mean ± SD. (E) Distance of the bacteria to the centroid of the host cell. Graph represents the median (8.74 µm) and interquartile range (6.51 – 11.05 µm). Values from n = 4990 cells from 7 independent experiments. (F) Distance of individual bacteria to the centroid of the host cell depending on its bacterial load. Graph represents the median and interquartile range. Kruskal-Wallis test and Dunn’s multiple comparisons test. (G) Area of cells depending on infection. Graph represents the median area (uninfected: 3031 µm^2^, infected: 4148 µm^2^) and interquartile range (unifected: 2308 – 4169 µm^2^, infected: 3062 – 5752 µm^2^). Values from n = 9469 uninfected cells and n = 1031 infected cells from 7 independent experiments. (H) Area of infected cells depending on its bacterial load. Graph represents the median and interquartile range. Kruskal-Wallis test and Dunn’s multiple comparisons test. (I) Nuclear area of cells depending on infection. Graph represents the median area (uninfected: 1551 µm^2^, infected: 1897 µm^2^) and interquartile range (unifected: 1209 - 1950 µm^2^, infected: 1486 - 2333 µm^2^). Values from n = 9469 uninfected cells and n = 1031 infected cells from 7 independent experiments. (J) Nuclear area of infected cells depending on its bacterial load. Graph represents the median and interquartile range. Kruskal-Wallis test and Dunn’s multiple comparisons test.

### Epithelial cells reduce *de novo* DNA and protein synthesis upon infection

In order to understand the physiological state of enlarged epithelial cells infected with *S. flexneri*, we monitored host DNA replication and protein translation using Click-chemistry. Using a 1 h pulse of the clickable thymidine analogue Edu, we specifically visualized and quantified *de novo* DNA synthesis in the population of epithelial cells with high-content microscopy (**Figure 2A, Figure 2-figure supplement 1**). We identified the subpopulation of cells in S-phase by their high level of Edu incorporation. Consistent with this, this subpopulation of cells was fully abrogated in the presence of the S-phase blocker Aphidicolin (Pedrali-Noy et al., 1980) (**Figure 2-figure supplement 2A, B**). Approximately one third of uninfected cells (34.45%) were found in S-phase (**Figure 2B, C**), in agreement with S-phase lasting around one third of the cell cycle (∼8h out of ∼22h). Infected cells presented a similar proportion of cells in S-phase (35.89%), indicating that *S. flexneri* infection did not block transition of cells from G1 to S at the timepoint tested. We then assessed the amount of *de novo* DNA synthesis incorporated in cells in S-phase during infection. Surprisingly, we observed a reduction of Edu incorporation in infected cells (**Figure 2D**), which was dependent on the bacterial burden (**Figure 2E**). These results indicate that *S. flexneri* infection slows down DNA synthesis of epithelial cells.

**Figure 2:**
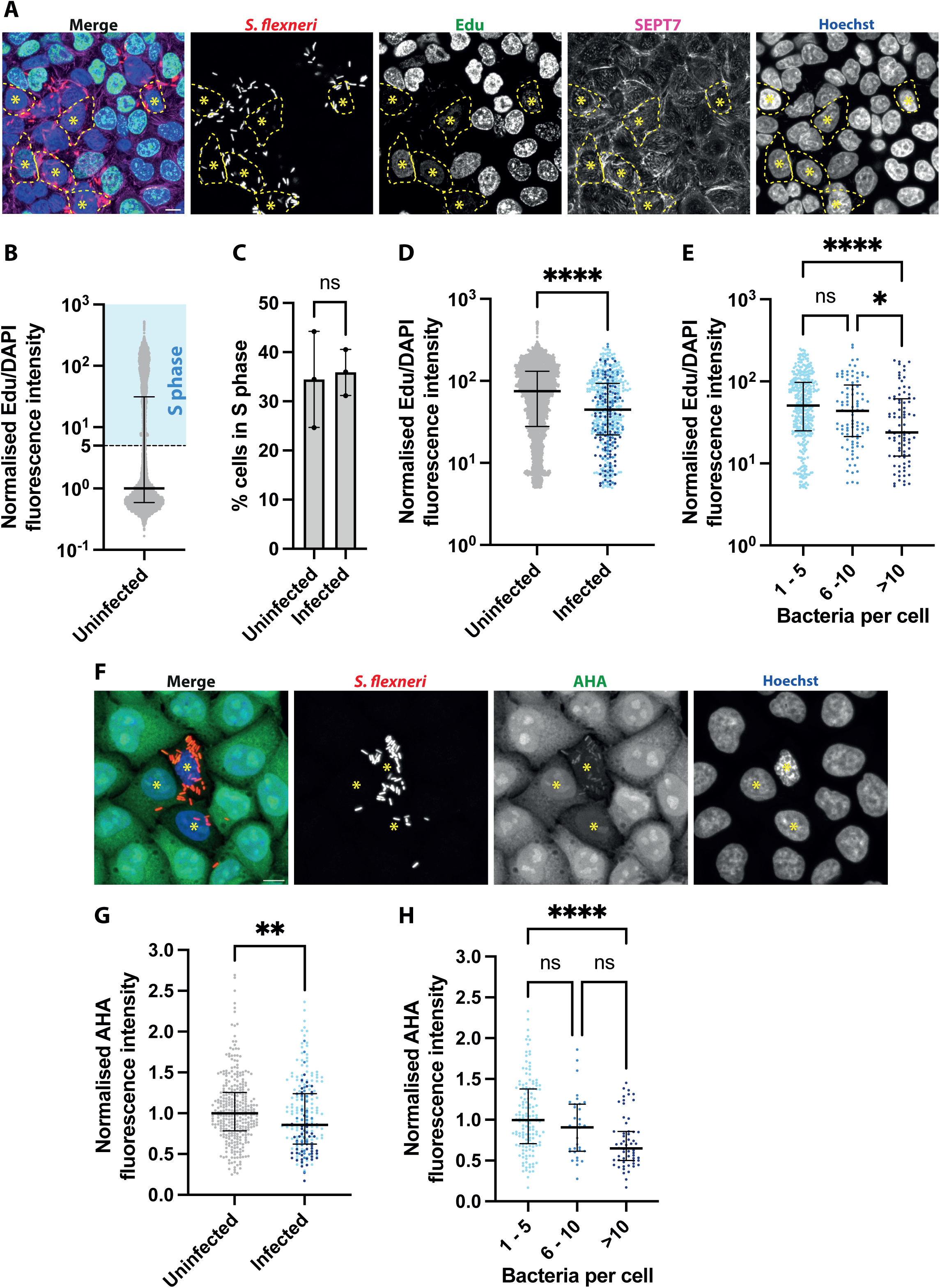
*S. flexneri* infection reduces host DNA synthesis and host protein synthesis. (A) Representative microscopy image of HeLa cells infected with *S. flexneri* expressing mCherry at 3 h.p.i, after 1h incubation with Click-IT Edu. Asterisks label the nucleus of infected cells with reduced Edu incorporation. Contour of selected cells is delimited with a dashed line. Scale bar, 10 µm. (B) Edu acquisition in uninfected cells is bimodal. Graph represents the normalised median of the ratio between the fluorescence intensity of Edu and Hoechst (1), and interquartile range (0.5924 – 31.22). Values from n = 8941 cells from 3 independent experiments. An arbitrary threshold of 5 is selected to define cells in S-phase. (C) Frequency of cells in S-phase (as defined in B) depending on infection. Graph represents mean ± SD (uninfected: 34.45% ± 9.78%, infected: 35.89% ± 4.69%). Student’s t-test. (D) Edu acquisition in cells in S-phase depending on infection. Graph representes the normalised median of the ratio between the fluorescence intensity of Edu and Hoechst (uninfected: 75.14, infected: 44.73), and interquartile range (uninfected: 27.89 – 131.0, infected: 22.05 – 93.36), from 3065 uninfected and 552 infected cells. Mann-Whitney U test. (E) Edu acquisition in cells in S-phase depending on bacterial load. Graph represents the median and interquartile range. Kruskal-Wallis test and Dunn’s multiple comparisons test. (F) Representative microscopy image of HeLa cells infected with *S. flexneri* expressing mCherry at 3 h.p.i, after 1h incubation with Click-IT AHA. Asterisks label the infected cells with reduced AHA incorporation. Scale bar, 10 µm. (G). AHA acquisition in host cells depending on infection. Graph representes the normalised median of AHA fluorescence intensity (uninfected: 1, infected: 0.86), and interquartile range (uninfected: 0.7850 – 2.694, infected: 0.6207 – 1.241), from 36 uninfected and 233 infected cells. Mann-Whitney U test. (H) AHA acquisition in infected cells depending on bacterial load. Graph represents the median and interquartile range. Kruskal-Wallis test and Dunn’s multiple comparisons test.

To assess protein translation, we used a 1h pulse of the clickable methionine analogue AHA, which specifically labels *de novo* synthesized proteins as AHA incorporation is reduced after treatment with the inhibitor of translational elongation Cycloheximide (Ennis & Lubin, 1964) (**Figure 2-figure supplement 1C, D**). During infection, we observed a reduction of protein translation in infected cells compared to non-infected cells (**Figure 2F,G**), which was also dependent on the bacterial burden (**Figure 2H**).

Collectively, the cell morphological rearrangements together with the reduction of *de novo* DNA and protein synthesis in infected epithelial cells suggests that *S. flexneri* induces an arrest in the host cell cycle.

### Design of a deep learning pipeline to automatically identify bacteria interacting with SEPT7

*S. flexneri* have been shown to interact with septins during infection. These interactions include septin ring-like structures surrounding phagocytic cups, autophagosomes, actin tails and protrusions, all being morphologically diverse structures that are normally scored by hand (Krokowski et al., 2018; Lobato-Márquez et al., 2023; Mostowy et al., 2010) . To study septin-associated bacteria using high-content microscopy, we designed a tailored analysis pipeline for the automatic and unbiased identification of SEPT7-bacteria interactions (**Figure 3**). SEPT7 is a core component of septin hexamers and octamers, and is historically used to identify septin structures (Mostowy et al., 2010). The analysis pipeline includes two deep learning models based on Convolutional Neural Networks (CNN), which are effective in solving classification tasks in computer vision (Krizhevsky et al., n.d.; Lecun et al., 2015; Nielsen, 2015). As septins can surround individual and also multiple bacteria simultaneously (**Figure 4-figure supplement 1A**), we focused on individual bacteria. For this, we trained the first classification model to discriminate between “single” bacteria and “clumps”. The second classification model was designed to assign those single bacteria into “SEPT7-positive structures” or alternatively “negatives”, according to the morphology of SEPT7 recruitment around them.

**Figure 3.**
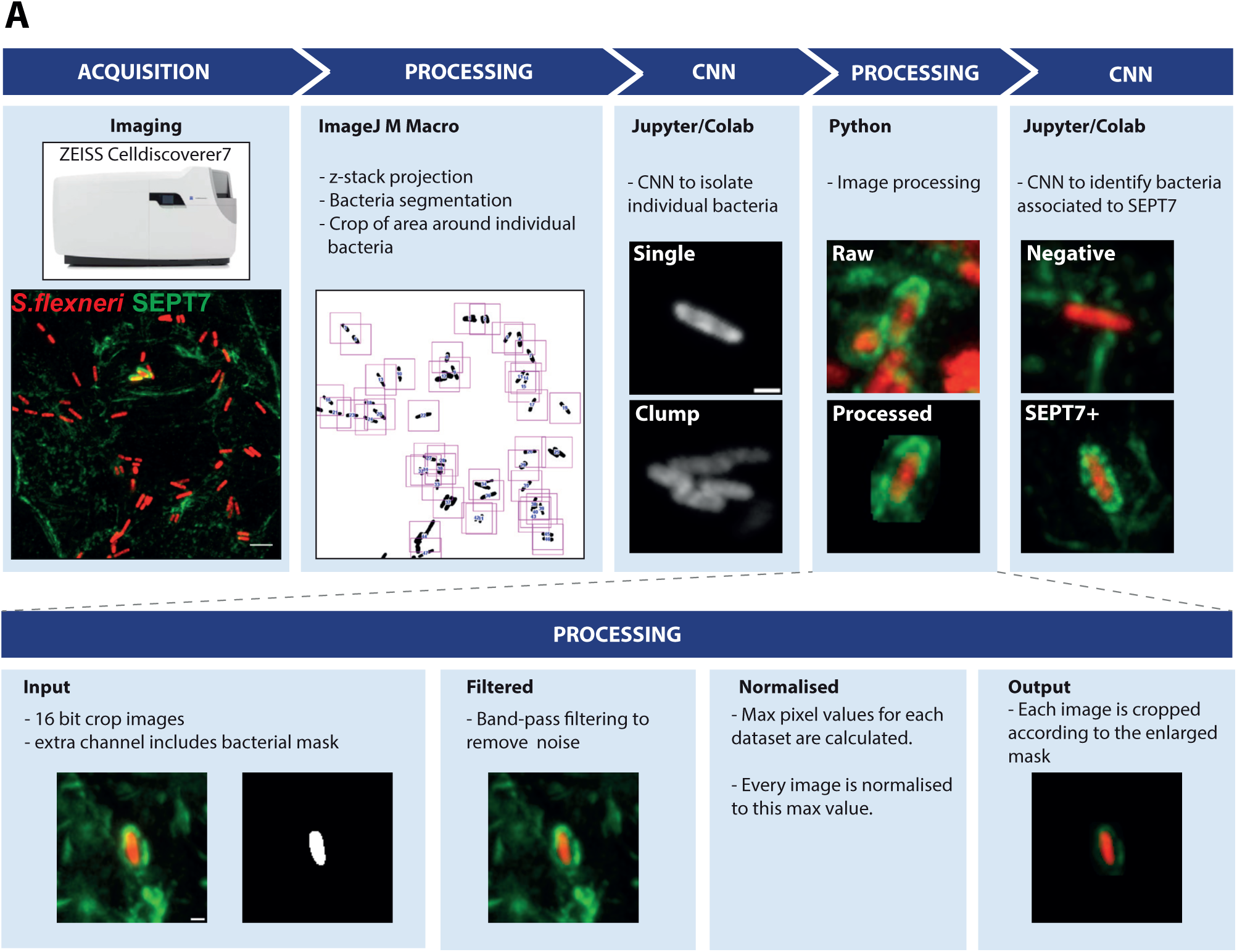
Microscopy pipeline to automatically identify SEPT7-*S. flexneri* interactions. (A) Diagram representing the steps involved in the imaging and analysis pipeline. After infection, fixation and staining, high number of images are automatically acquired using the high-content microscope. Scale bar, 10 µm. These are then processed to segment bacteria based on their fluorescence, so that a square field containing the bacteria in the centre is cropped and saved for subsequent analysis. A deep learning model based on CNN is applied using Jupiter notebooks and the Python Keras library hosted on Colab to sort individual, isolated bacteria. Several processing steps are applied using Python to remove noise (band-pass filtering by difference of Gaussian and mean filters), normalise the data across datasets for comparison, and remove signal away from the bacteria which is irrelevant for the identification of SEPT7 assemblies. Scale bar, 1 µm. Finally, a similar second deep learning model based on CNN is applied to identify SEPT7 interacting with bacteria.

The first classification model (discerning “single” versus “clumped” bacteria) together with its training and performance metrics is summarized in **Figure 4**. A large dataset of annotated bacteria (**Figure 4A, B, Figure 4-figure supplement 1B**) was used to train a Sequential model with a typical CNN architecture that was experimentally tested and optimised accordingly for best results (**Figure 4C**). This model provided a good fit of the training data (high accuracy, low loss), with similar predictive power for the validation dataset (**Figure 4D, E**). Performance of the model was further assessed with a confusion matrix where a separate annotated test dataset was challenged and correct number of predictions scored, showing that 96% of SEPT7 structures and 83% of clumps were correctly identified (**Figure 4F**). Finally, effective performance of the model was ensured with high values achieved in additional metrics testing for type I and II errors (precision, recall, F1-score) (**Figure 4G**).

**Figure 4.**
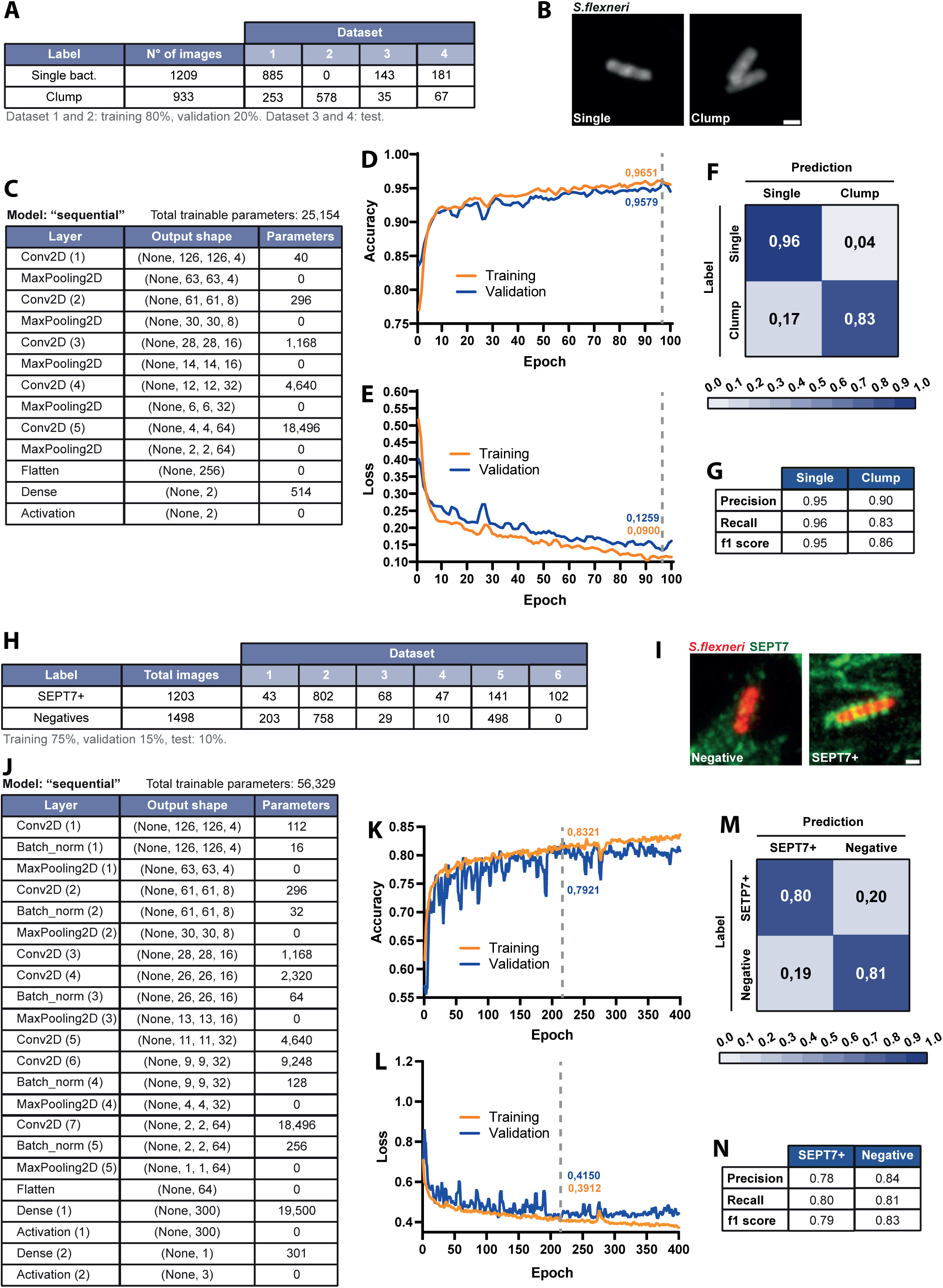
Training of a deep learning algorithm to identify single isolated bacteria cells and SEPT7 associated *S. flexneri*. (A) Table summarizing the annotated dataset used for the training process. The labels of the two classification classes used were “Single bacteria” and “Clump”. The annotated images were randomly split into two groups, one used as training dataset (comprising 80% of the images) and another used as validation dataset (with the remaining 20%). A separate annotated dataset was used for testing. (B) Representative Airyscan images of bacteria used for each class. 2D max projections of Z-stacks were used for training. Scale bar, 1 µm. (C) Architecture of the deep learning algorithm used for training. Each raw describes the characteristics of the sequential transformation steps applied. Conv2D stands for 2D Convolution. (D, E) Accuracy and Loss as metrics to represent the training process over subsequent epochs (entire passing of the training data through the algorithm). Accuracy increases and loss decreases for both training and validation datasets, indicating a good fit of the model to the data. Vertical dashed grey line indicates Early Stopping, or epoch value for the minimum validation Loss. (F) Confusion matrix performed on an annotated test dataset not used previously for the training or validation process, indicating the percentage of predictions that were correct or wrong for each class. (G) Precision, recall and f1 score as metrics that summarize the performance of the model or classifier. (H) Table summarizing the annotated dataset used for the training process. The labels of the two classification classes used were “Septin” and “Negative”. The annotated images were randomly split into three groups, one used as training dataset (comprising 75% of the images), another used as validation dataset (with 15%) and a last one used as test dataset (with 10%). Due to the data being very imbalanced (15% natural frequency of SEPT7 associated bacteria), the images comprised in the Negative class were under-sampled as indicated in the table. (I) Representative Airyscan images of bacteria used for each class. 2D max projections of Z-stacks were used for training. Scale bar, 1 µm. (J) Architecture of the deep learning algorithm used for training. Each raw describes the characteristics of the sequential transformation steps applied. Conv2D stands for 2D Convolution, Batch_norm stands for Batch normalisation. (K, L) Accuracy and Loss as metrics to represent the training process over subsequent epochs. Accuracy increases and loss decreases for both training and validation datasets, indicating a good fit of the model to the data. Vertical dashed grey line indicates Early Stopping, or epoch value for the minimum validation Loss. (M) Confusion matrix performed on the test dataset not used previously for the training or validation process, indicating the percentage of predictions that were correct or wrong for each class. (N) Precision, recall and f1 score as metrics that summarize the performance of the model or classifier.

The second classification task (identification of SEPT7 “positive” vs “negative” bacteria) was more challenging due to their intrinsic heterogeneity plus the presence of ubiquitious SEPT7 filaments that populate the vicinity of bacteria (but do not associate with bacteria per se). To address these challenges, we increased the amount of data and complexity of the second classification model (**Figure 4H-J, Figure 4-figure supplement 1C**) compared to the first (**Figure 4A-C**). This included more than 1200 images of *S. flexneri* associated to SEPT7, both from infections with wild-type bacteria or expressing AfaI to increase bacterial invasion. The training process achieved a high accuracy and low loss for the training and validation dataset (**Figure 4K, L**). Correct prediction of manually annotated SEPT7 positive structures was 80% as shown in the confusion matrix (**Figure 4M**), with performance metrics Precision, Recall and F1 score also around 0.8 (**Figure 4N**).

### Characterization of the morphological heterogeneity of SEPT7 recruitment to *S. flexneri*

The training of the classification model to identify bacteria associated to SEPT7 required a large dataset of images. We observed diverse morphological patterns of SEPT7 signal around bacteria (**Figure 5 A-C**). We differentiated five frequent subcategories from a total of 855 cases. The first corresponds to most cases (66.7%), where SEPT7 forms rings that entrap bacteria transversally (**Figure 5A, B, a**). We also observed two categories where SEPT7 surrounds bacteria more homogeneously either in a tight (18.2%) or loose association (6.1%) (**Figure 5A, B, b-c**). Finally, we observed partial SEPT7 recruitment, with only one (5.0%) or two (4.0%) bacterial poles being targeted (**Figure 5A, B, d-e**). Given this diversity, we combined the entire dataset to summarise the probability of SEPT7 interactions with bacteria (**Figure 5-figure supplement 1**). As bacteria are found in any orientation inside epithelial cells, we included only those instances where the z-projection of the bacterial axis measured more than 2.5µm, ensuring they were positioned horizontally when imaged. Representations shown in **Figure 5-figure supplement 1A, C** and graphs in **Figure 5-figure supplement 1B, D** can be understood as a probability map of the area of influence of SEPT7 when associating with intracellular *S. flexneri*.

**Figure 5.**
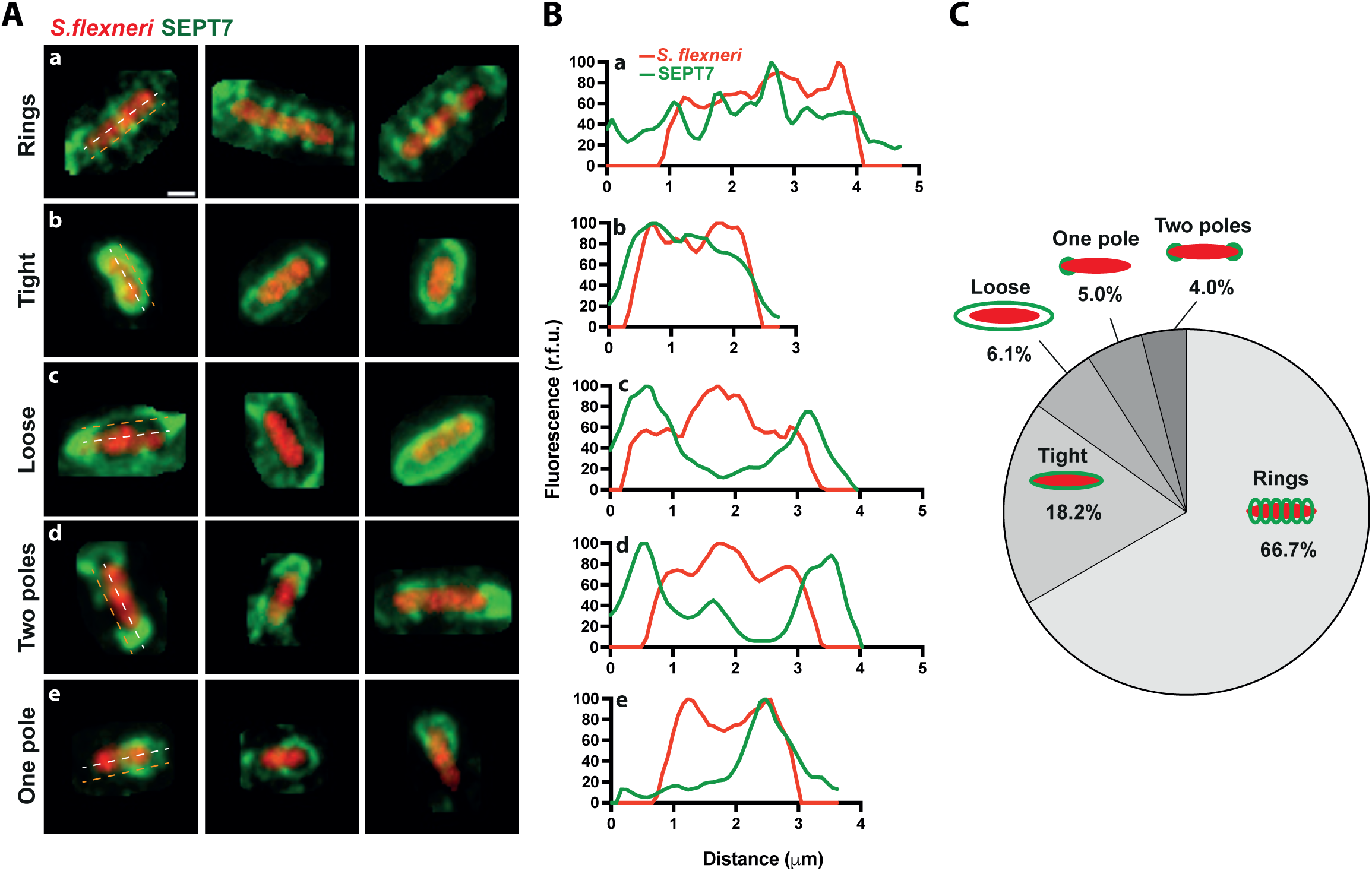
Septin assemblies associated to *S. flexneri* are heterogeneous. (A) Selected examples of the training dataset to showcase the diverse morphology of septin assemblies associated to bacteria. Frequent morphologies include (a) ring like structures around the bacteria, (b) smooth tight recruitment around the entire bacterial surface, (c) loose structures more distant from the bacterial surface, (d) septin recruitment to both bacterial poles, (e) septin recruitment to a single bacterial pole. Scale bar, 1 µm. (B) Intensity profiles of the cytosolic bacterial mCherry and the SEPT7 recruitment, as indicated in the white and orange dashed lines, respectively. (C) Pie chart representing the relative frequency of 855 *S. flexneri*-SEPT7 associations depicted in A.

### SEPT7 is recruited to *S. flexneri* with increased T3SS activity

Septins have previously been shown to recognise growing bacterial cells *in vitro* and during infection of HeLa cells, using pharmacologic and genetic manipulation (Krokowski et al., 2018; Lobato-Márquez et al., 2021). To test the metabolic state of SEPT7-associated *S. flexneri* at 3h 40 min post infection (hpi) we used our high-content imaging and analysis pipeline (**Figure 6**). We measured *S. flexneri de novo* synthesis of DNA and proteins using a 1 h pulse Click-IT Edu and Click-IT AHA. Only active bacteria incorporated these analogues, as the antibiotic Nalidixic acid (inhibitor of the bacterial DNA gyrase (Sugino et al., 1977)) and Rifampicin (inhibitor of the bacterial RNA polymerase (Ezekiel & Hutchins, 1968)) significally reduced their incorporation, respectively (**Figure 6-figure supplement 1A-D, Figure 6-figure supplement 1E,F)**. During infection, bacteria associated with SEPT7 showed heterogeneus levels of DNA synthesis and protein translation similar to SEPT7-negative bacteria (**Figure 6A-H**).

**Figure 6.**
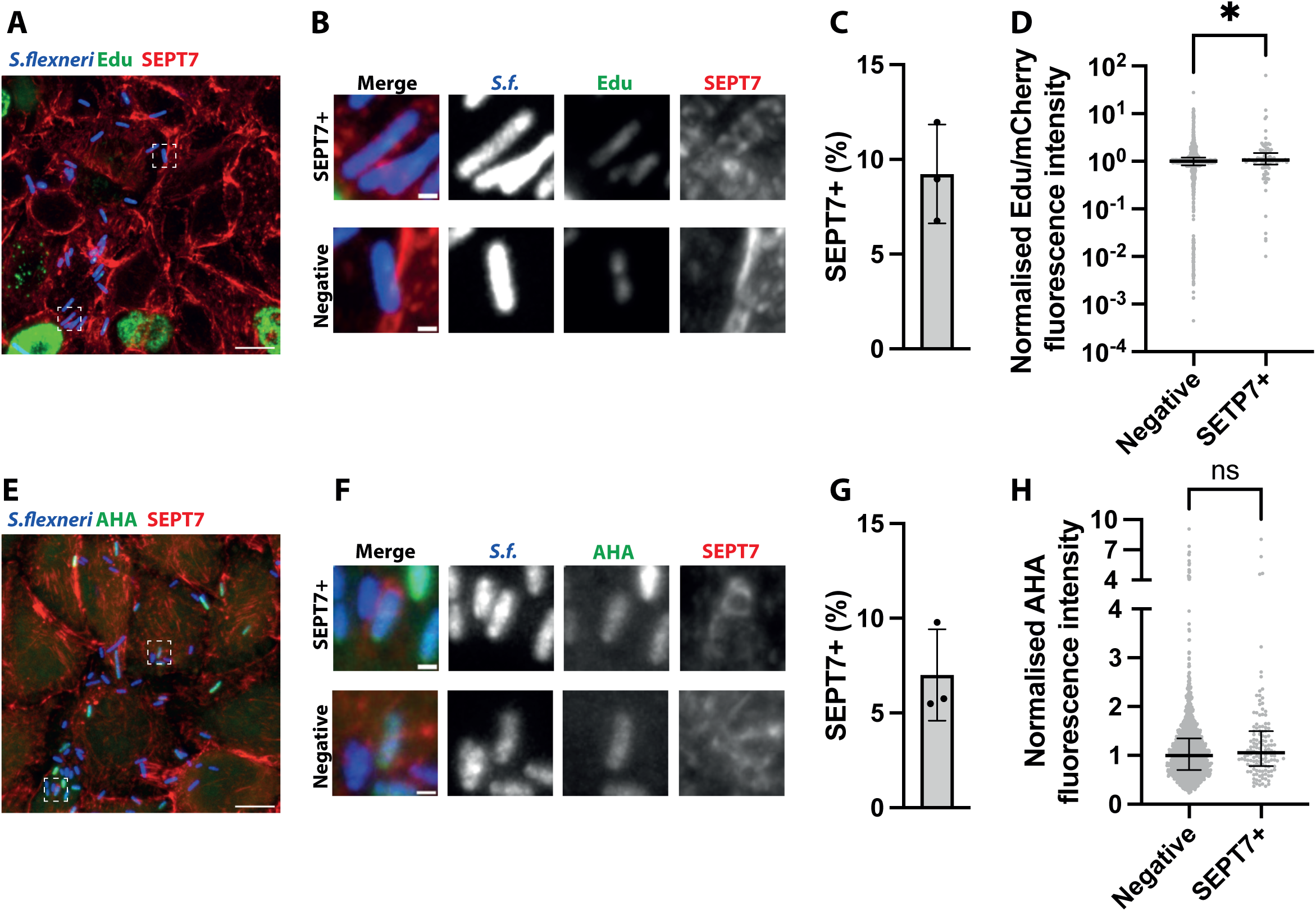
*S. flexneri* associated with SEPT7 are metabolically active. (A) Representative microscopy image of HeLa cells infected with *S. flexneri* expressing mCherry at 3 h.p.i, after 1h incubation with Click-IT Edu. Scale bar 10 µm. (B) Enlarged view of insets in A, highlighting a *S. flexneri* associated or not to SEPT7 structures. Scale bar, 1 µm. (C) Frequency of the number of SEPT7-associated bacteria identified using deep learning from experiments in A, from n= 1024 from 3 independent replicates. Graph represents mean ± SD (9.23% ± 2.62%). (D) Normalised ratio between Edu signal and mCherry in individual bacteria. Graph representes the median (negatives: 1, septin associated: 1.059), and interquartile range (negatives: 0.8185 – 1.207, septin associated: 0.8573 – 1.487). Mann-Whitney U test. (E) Representative microscopy image of HeLa cells infected with *S. flexneri* expressing mCherry at 3 h.p.i, after 1h incubation with Click-IT AHA. Scale bar 10 µm. (F) Enlarged view of insets in E, highlighting a *S. flexneri* associated or not to SEPT7 assemblies. Scale bar, 1 µm. (G) Frequency of the number of SEPT7-associated bacteria identified using deep learning from experiments in E, from n = 1872 bacteria from 3 independent replicates. Graph represents mean ± SD (7.01% ± 2.41%). (D) Normalised AHA signal in individual bacteria in A, B. Graph representes the median (negatives: 1, SEPT7 associated: 1.055), and interquartile range (negatives: 0.6986 – 1.345, SEPT7 associated: 0.7837 – 1.497). Mann-Whitney U test.

*S. flexneri* pathogenicity relies on its T3SS. To understand effector secretion at the individual bacterial level, we used *S. flexneri* expressing the Transcription-based Secretion Activity Reporter (TSAR) (Campbell-Valois et al., 2014). This reporter comprises a short lived fast-maturing superfolder GFP (GFPmsf2) that is expressed in conditions where the T3SS is active, such as in broth upon addition of congo red (**Figure 7-figure supplement 1A, B**) (Parsot et al., 1995) and during infection (**Figure 7**). We observed heterogeneity in TSAR signal (**Figure 7A,B**). T3SS activation was higher in bacteria found in cells with a low bacterial burden (less than 10 intracellular bacteria) (**Figure 7A,C**). This suggests the highest level of activation occurs in bacterial pioneers that colonise neighbouring cells, in agreement with previous reports (Campbell-Valois et al., 2014). We then assessed T3SS activation in *S. flexneri* associated with SEPT7 using our deep learning analysis pipeline (**Figure 7D-H**). We detected a significant increase in TSAR-positive bacteria in SEPT7 associated *S. flexneri* compared to SEPT7-negative bacteria (**Figure 7G,H**). These include SEPT7-associated protrusions and cage-like structures (**Figure 7E**, **Figure 7-figure supplement 1C**). These data underscore the antimicrobial role of septins in associating with actively secreting bacteria.

**Figure 7.**
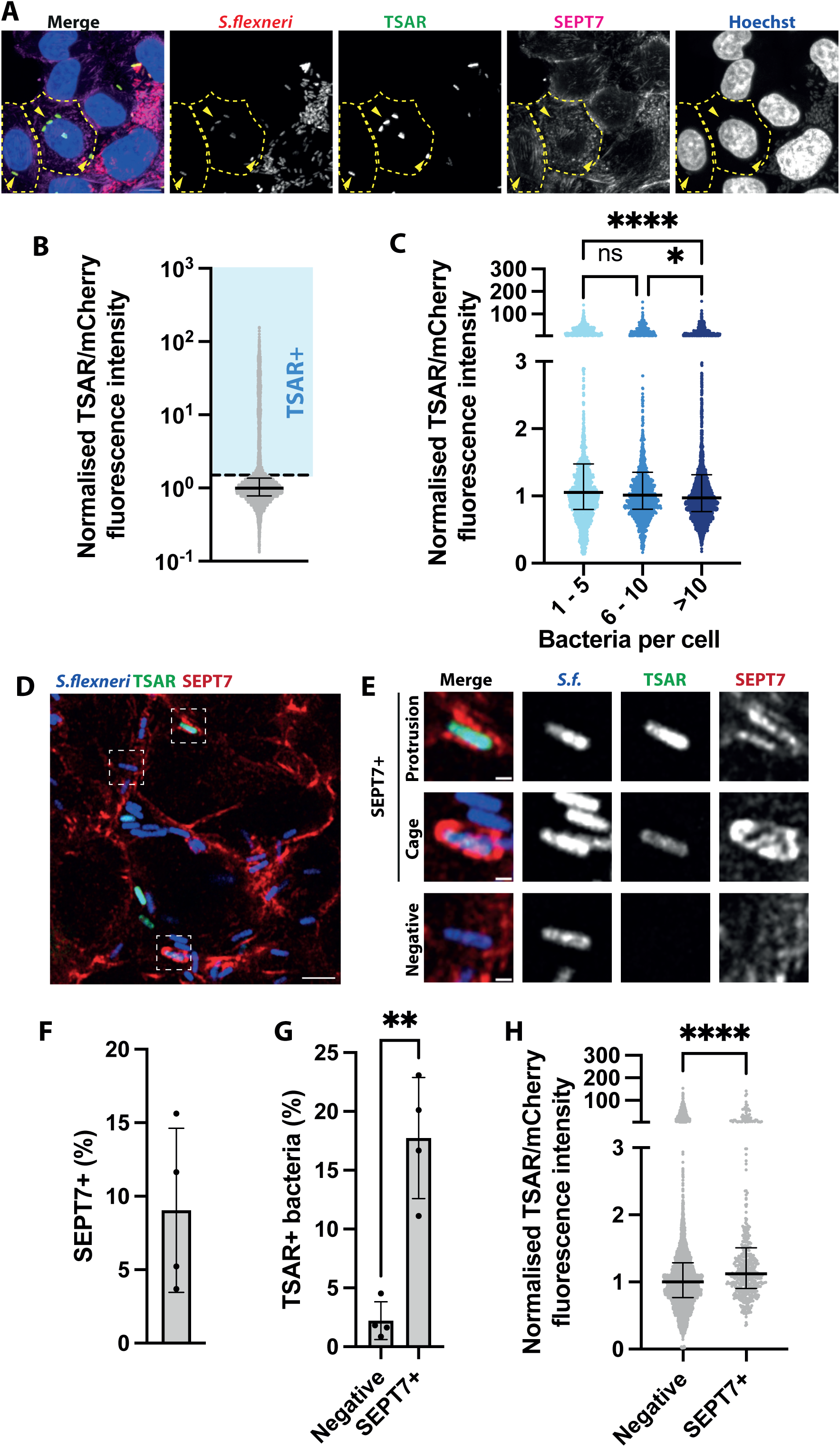
*S. flexneri* associated with SEPT7 have an active T3SS. (A) Representative microscopy image of HeLa cells infected with *S. flexneri* expressing the TSAR reporter at 3 h.p.i. Scale bar 10 µm. Arroheads highlight TSAR+ bacteria. Selected infected cells contours are delimited with dashed lines. (B) Distribution of TSAR signal intensity in individual bacteria. Graph represents the normalised median of the ratio between TSAR and mCherry (1), and interquartile range (0.7827 – 1.369). Values from n = 3654 bacterial cells from 3 independent experiments. An arbitrary threshold of 1.5 is selected to define TSAR+ *S. flexneri*. (C) Intensity of TSAR signal in individual bacteria depending on infection load of host cells. Graph represents the normalised median and interquartile range. Kruskal-Wallis test and Dunn’s multiple comparisons test. (D) Representative microscopy image of HeLa cells infected with *S. flexneri* expressing the TSAR reporter at 3 h.p.i. Scale bar 10 µm. (E) Enlarged view of insets in D, highlighting a *S. flexneri* associated or not to SEPT7 assemblies. Scale bar, 1 µm. (F) Frequency of the number of SEPT7-associated bacteria identified using deep learning from experiments in F, from n= 5030 bacteria from 4 independent replicates. Graph represents mean ± SD (9.044% ± 5.582%). (G) Percentage of TSAR+ *S. flexneri* depending on their association to SEPT7. Graph represents mean ± SD (negatives: 2.214 ± 1.603, septin associated: 17.74 ± 5.138). Student’s t-test. (H) Normalised ratio between TSAR signal and mCherry in individual bacteria. Graph representes the median (negatives: 1, SEPT7 associated: 1.122), and interquartile range (negatives: 0.7674 – 1.286, SEPT7 associated: 0.9047 – 1.511). Mann-Whitney U test.

## Discussion

Unravelling the intrinsic heterogeneity of host and bacterial cells underlying the infection process is paramount to design effective therapeutic strategies. High-content and high-throughput image analysis are ideal tools to capture this heterogeneity when coupled to automatic phenotypic image analysis (Eliceiri et al., 2012; Mattiazzi Usaj et al., 2016; Pegoraro & Misteli, 2017). Here we use high-content high-resolution microscopy and automated image analysis to quantitatively dissect the heterogeneity of *S. flexneri* in epithelial cells at the single-cell and population level at unprecedented subcellular detail.

We discover that *S. flexneri* induces important morphological and physiological changes in epithelial cells after 3h40min post infection. These include a marked decrease of DNA synthesis in cells in S-phase. Consistent with the ability of bacteria to manipulate the host cell cycle, *S. flexneri* has been shown to induce genotoxic stress (Bergounioux et al., 2012) and block the G2/M phase transition (Iwai et al., 2007; Wang et al., 2019). We also observed a collective shutdown of protein expression, which happened in a bacterial-burden dependent manner. A recent report has shown that the OspC family of *S. flexneri* effectors ADP-riboxanates the host translation initiation factor 3, leading to translational arrest and the formation of stress granules (Zhang et al., 2024). Collectively, these observations suggest that *S. flexneri* infection induces host cell stress and arrests the progression of the cell cycle. In the natural *S. flexneri* niche of the human gut, such an arrest would prevent maturation and shedding of infected epithelial cells, thereby promoting efficient gut colonisation (Iwai et al., 2007).

To understand how bacteria are targeted by cell-autonomus immune defences, such as septin cage entrapment, it is necessary to investigate phenotypic characteristics of targeted bacteria and the heterogeneous recruitment of cell-autonomous immune factors. While traditional image segmentation tools fail to enable this task, rapidly evolving developments in the field of computer vision are democratising the use of deep learning and CNNs for multiple applications (Berg et al., 2019; Hung et al., 2020; Stringer et al., 2020; von Chamier et al., 2021; Weigert et al., 2020) including infection biology (Davidson et al., 2021; Fisch et al., 2019b, 2021; Touquet et al., 2018). However, the analysis of complex datasets still requires a tailored approach, as shown here for bacterial interactions with the septin cytoskeleton (which is ubiquitious in the cell and forms diverse assemblies whose precise identification requires high resolution). We successfully implement a CNN based approach for the automated classification of bacteria associated with SEPT7 assemblies using z-stacks projections, enabling the analysis of thousands of bacterial cells while saving time and preventing human bias. Building upon this work, future studies will improve use of 3D data for CNN re-training instead of z-stack projections, to allow more precise identification of *S. flexneri*-septin interactions, particularly when bacteria are placed perpendicular to the imaging plane.

A large dataset of images depicting *S. flexneri* – SEPT7 interactions was acquired to train the deep learning algorithm and to characterize the diversity of these structures. We observe that the most frequent septin assembly is in the form of rings transversally surrounding the bacterial surface. We also detected alternative morphologies, including homogeneous SEPT7 labelling surrounding the bacterial surface (tight vs loose), as well as the discrete recruitment of SEPT7 to one or two bacterial poles. While previous work has shown that septin cage assembly around *S. flexneri* can start with the discrete recruitment of septins to one or two bacterial poles (Krokowski et al., 2018), it is not yet known if the different septin assemblies we describe here correspond to different stages or fates of the same structure or whether it marks separate biological events. To fully address this question, follow-up studies should include timelapse analysis of septin cage formation. We use deep learning analysis to interrogate intracellular bacteria and show that *S. flexneri* associated with SEPT7 present heterogenous DNA and protein synthesis. In addition, we show that bacteria associated with SEPT7 have increased T3SS activity (as compared to bacteria not associated with SEPT7). T3SS activity has been shown to be higher in motile bacteria (Campbell-Valois et al., 2014) which reinforces the antimicrobial role of septins in counteracting actin tail motility. Altogether, our deep learning tool to analyse septin-bacteria interactions opens the door to future studies on interactions between bacteria and components of cell-autonomous immunity.

## Concluding remarks

Heterogeneity in a controlled lab environment is only a small representation of the broader heterogeneity underlying the clinical infection process, considering the genetic background of the patient, interactions with microbiota, etc. In the future, our pipeline can be applied to investigate septin-bacteria interactions among different *Shigella* spp. or clinically relevant strains, as well as other intracellular bacteria (eg. *M. marinum, Staphylococcus aureus*, *Pseudomonas aeruginosa*). In addition, our pipeline can be adapted to identify and score other cytoskeletal structures (such as actin tails) or membrane organelles (such as autophagosomes) associated with intracellular bacteria. Ultimately, we envision that our automated microscopy workflows will help to screen pharmacologic or genetic libraries in a multiparametric manner and understand their impact on host and bacterial cells at the single-cell and population level. In this way, novel host and bacterial factors underlying heterogeneity and bacterial pathogenesis could be identified, leading to transformative therapeutic strategies to combat bacterial infection in humans.

## Materials and methods

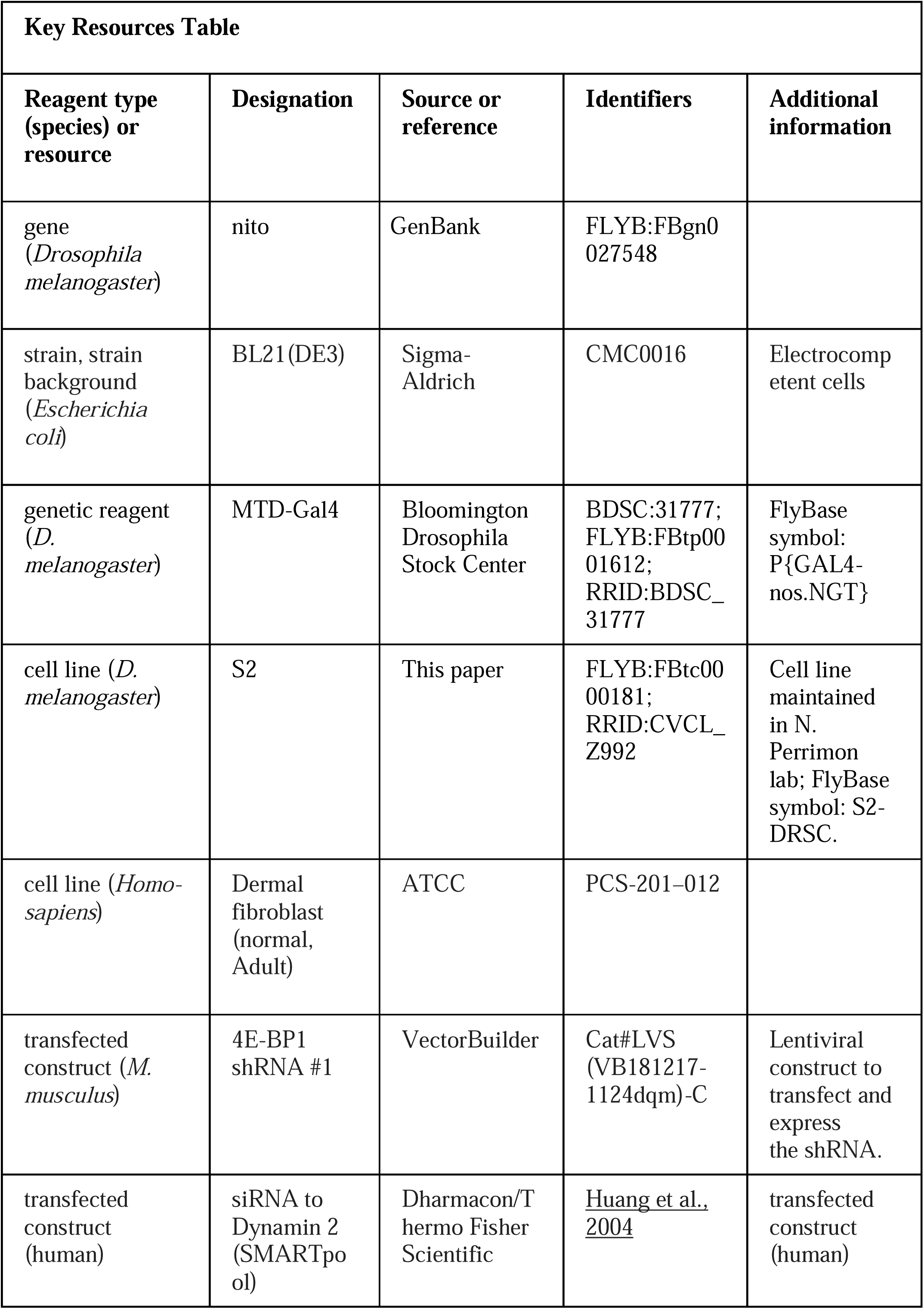
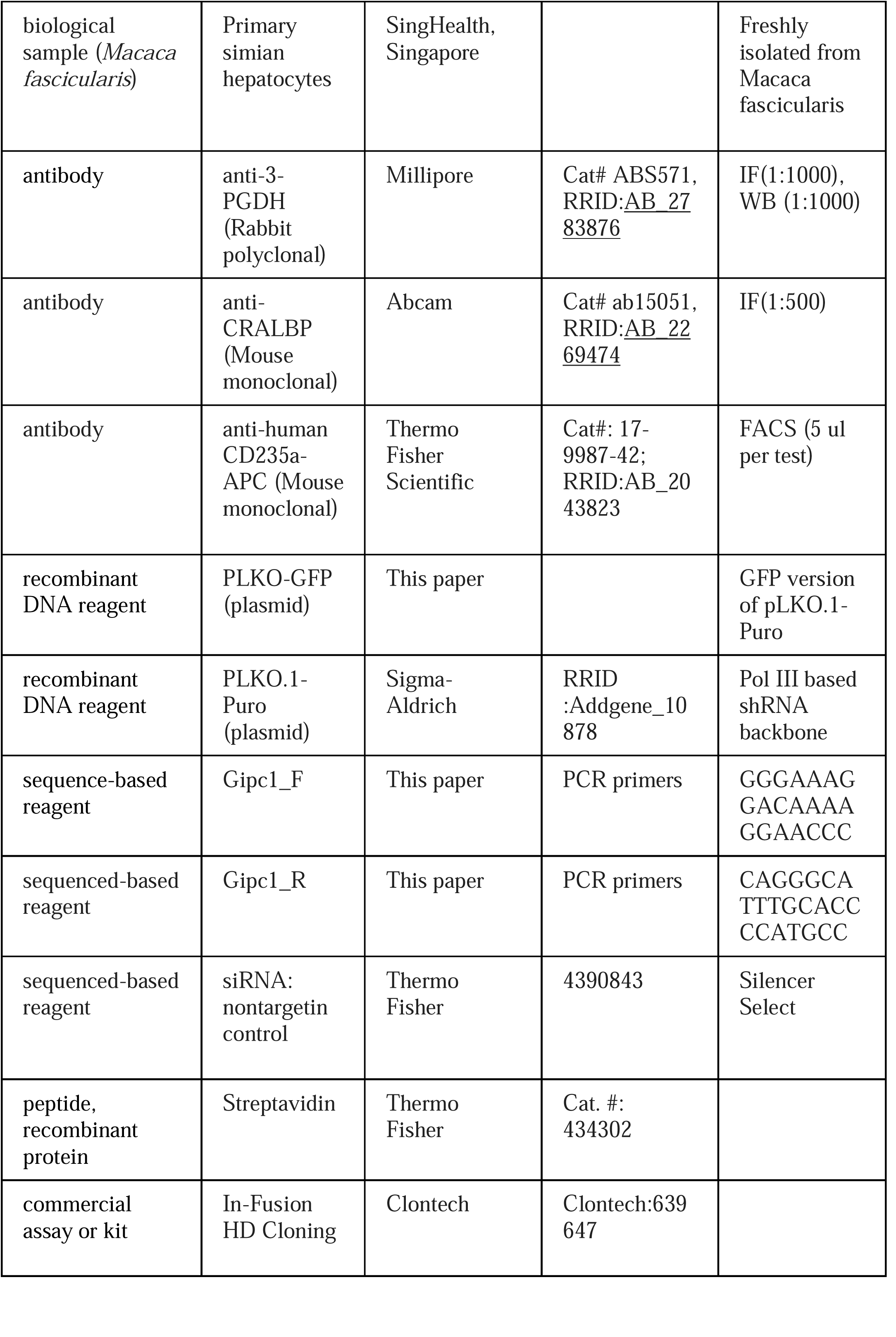
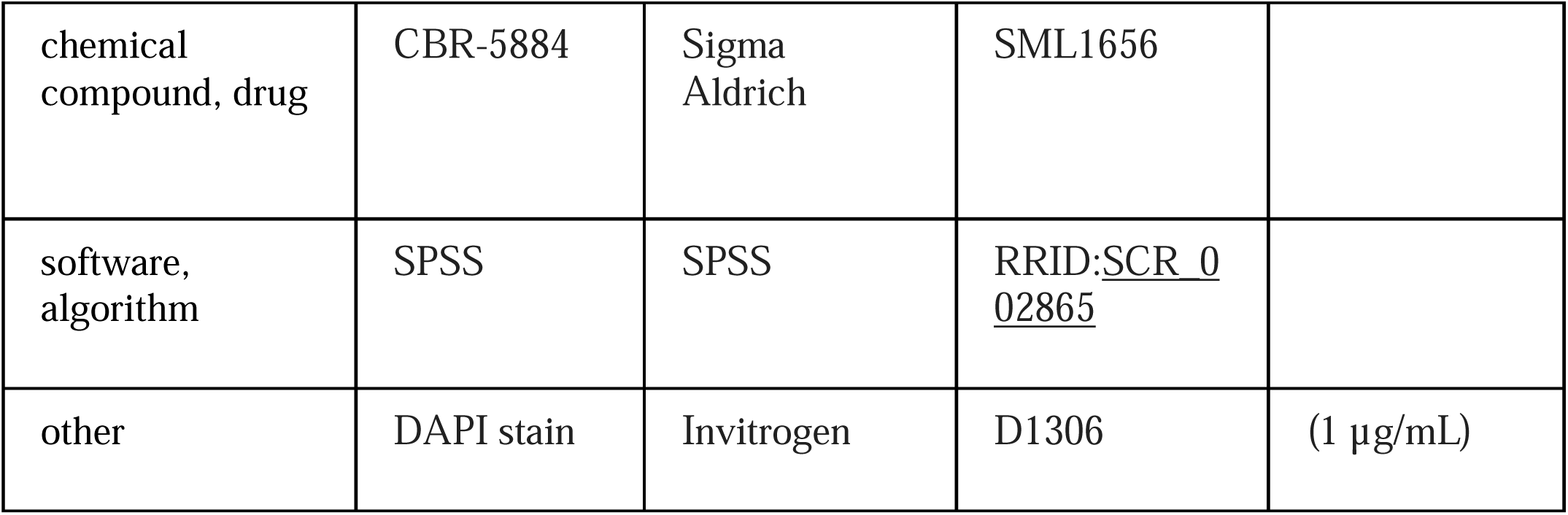

### Reagents

The following antibodies were used: rabbit anti-SEPT7 (1:100 dilution, #18991, IBL), Alexa-488-conjugated anti-rabbit antibody (1:500, #10729174, ThermoFisher Scientific), Alexa-647-conjugated anti-rabbit antibody (1:500, #A27040, ThermoFisher Scientific). The following dyes were used: Alexa-647-conjugated phalloidin (1:500, #10656353, ThermoFisher Scientific), Hoechst (1:500, #H3570, ThermoFisher Scientific), 0.01% (w/v) congo red (#C6767, Sigma-Aldrich). The following drugs were used: 50 μg/mL aphidicolin (#A0781 Sigma-Aldrich), 50 μg/mL cycloheximide (#2112 Cell Signalling), 2 μg/mL nalidixic acid (#N8878 Sigma-Aldrich), and 100 μg/ml rifampicin (#R3501 Sigma-Aldrich).

### Bacterial strains and culture conditions

The bacterial strains and plasmids described in this study are listed in **Sup. Table 1**. *Shigella flexneri* 5a str. M90T mCherry was used throughout the manuscript unless indicated otherwise unless indicated otherwise. *S. flexneri* strains were grown in trypticase soy broth (TCS) agar containing congo red to select for red colonies, indicative of a functional T3SS. Conical polypropylene tubes (#CLS430828, Corning) containing 5 ml of TCS were inoculated with individual red colonies of *S.flexneri* and were grown ∼16 h at 37 °C with shaking at 200 rpm. The following day, bacterial cultures were diluted in fresh prewarmed TCS (1:50 or 1:100 v/v), and cultured until an optical density (OD, measured at 600 nm) of 0.6.

### Cloning

Primers used in this study were designed using NEBuilder Assembly Tool (https://nebuilder.neb.com/#!/). To produce *S. flexneri* expressing AfaI and GFP, the pACAfaI plasmid was constructed (plasmids characteristics and primer sequences are located in **Supplementary Table 1**). Briefly, the *afaI* operon together with the *amp^R^* gene was amplified from *S.flexneri afaI* strain using the primers *afaI-operon-fw* and *afaI-operon-rv*. The *ori p15A* was amplified from pAC-P_lac_ using the primers *p15A-ori-fw* and *p15A-ori-fw.* Both DNA fragments were assembled using the Gibson assembly to generate the plasmid pAC-AfaI-AmpR. In a second step, the *AmpR* gene from pAC-AfaI-Amp was substituted by the *CmR* gene and generate pAC-AfaI-Cm. This was achieved by amplification of the *CmR* gene from pNeae2 using the primers *CmR-fw CmR-rv* and amplification of the backbone of pAC-AfaI-Amp with primers *afaI-p15A-fw, afaI-p15A-rv*, followed by Gibson assembly. Gibson assembly was performed at 50 °C for 30 min using the HiFi DNA Assembly Master Mix (#E2621L, New England Biolabs). Resulting plasmid pAC-AfaI-Cm was transformed into *S. flexneri* GFP by electroporation.

### Cell lines

HeLa (ATCC CCL-2) cells were grown at 37 °C and 5% CO_2_ in Dulbecco’s modified Eagle medium (DMEM, GIBCO) supplemented with 10% fetal bovine serum (FBS, Sigma-Aldrich). Cells lines were tested for mycoplasma infection and tested negative.

### Infection of human cells

Hela cells were seeded in 96-well microplates (PhenoPlates, Perkin Elmer) at a confluency of 10^4^ cells/well 2 days prior to infection. Alternatively, 9 × 10^4^ HeLa cells were seeded in 6-well plates (Thermo Scientific) containing 22 × 22 mm glass coverslips 2 days before the infection. Bacterial cultures were grown as described and cell cultures were infected with *S. flexneri* strains as described previously (Mostowy et al., 2010). Briefly, Hela cells were infected with *S. flexneri* by spin-inoculation at 110 g for 10 min at a multiplicity of infection (MOI, bacteria:cell) of 100:1. Then, microplates were placed at 37 °C and 5% CO_2_ for 30 min. Infected cultures were washed 3 × with phosphate-buffered saline (PBS) pH 7.4 and incubated with fresh DMEM containing 10% FBS and 50 mg/mL gentamicin at 37 °C and 5% CO_2_ for 3 or 4 h. In order to obtain more instances of *S. flexneri* associated to SEPT7 in Figure 4H, one of the datasets was obtained after infection with *S. flexneri* expressing the adhesin AfaI (Labigne-Roussel & Falkow, 1988) and GFP to increase host cell invasion at a MOI of 10 in the absence of spin inoculation.

### Click chemistry

To label *de novo* DNA and protein synthesis, Click-iT™ EdU Cell Proliferation Kit for Imaging (#C10337 Thermofisher Scientific) and Click-iT™ AHA Protein Synthesis HCS Assay (#C10289 Thermofisher Scientific) were used, respectively, according to the manufacturer instructions. In the case of AHA, cells were cultured in glutamine, cystine and methionine-free DMEM (#21013024 Thermofisher Scientific) for 1 h prior to the treatment. 10μM EdU and 25μM AHA were supplemented to live bacterial or mammalian cell cultures for 1h. Cells were fixed for immunofluorescence as indicated below. Click azide/alkyne reactions were performed after permeabilization and prior to antibody staining as indicated below, according to the manufacturer’s instructions.

### Immunofluorescence and fluorescence microscopy

Bacteria, coverslips or 96-well microplates containing adherent infected or uninfected cells were washed three times with PBS pH 7.4 and fixed 15 min in 4% paraformaldehyde (in PBS) at RT. Fixed cells were washed three times with PBS pH 7.4 and subsequently permeabilized 5min with 0.1% Triton X-100 (in PBS). Cells were then washed three times in PBS and incubated 1h with primary antibodies diluted in PBS supplemented with 0.1% Triton X-100 and 1% bovine serum albumin. Cells were then washed three times in PBS and incubated 45 min with anti-rabbit secondary antibodies diluted 0.1% Triton X-100 (in PBS), and Hoestch and Alexa-conjugated phalloidin where indicated. 96-well microplates were washed three times in PBS and preserved in 0.01% sodium azide (#10338380 Thermo Scientific) in PBS pH 7.4. Stained bacteria cultures and coverslips were washed three times with PBS pH 7.4 mounted on glass slides with ProLong™ Diamond Antifade Mountant (#P36970I Invitrogen). Fluorescence microscopy on stained bacteria was performed using a using a 63×/1.4 C-Plan Apo oil immersion lens on a Zeiss LSM 880 confocal microscope driven by ZEN Blacksoftware (v2.3). Microscopy images were obtained using z-stack image series taking 8–16 slices. Fluorescence microscopy on infected or uninfected cells was performed using a ZEISS Plan-APOCHROMAT 20× / 0.95 Autocorr Objective or a ZEISS Plan-APOCHROMAT 50×/1.2 water immersion lens coupled to a 0.5x tubelens on a Zeiss CellDiscoverer 7 with Airyscan detectors driven by ZEN Blue software (v3.5). Microscopy images were obtained using z-stack image series taking 32 slices. Confocal images were processed using Airyscan processing (Weinerfilter) using “Auto Filter” and “3D Processing” options.

### Flow cytometry

5 x 10^4^ individual bacterial cells were analysed using flow cytometry with an LSRII flow cytometer (BD Biosciences). The data was analysed using FlowJo software, version 10.7.1. The median fluorescence of the bacterial population was determined and plotted for each staining.

### Analysis pipelines

Number of bacteria per cell, as well as fluorescence of bacteria, nuclei and mammalian cells was analysed using CellProfiler (v.4.0.7) (Carpenter et al., 2006). Custom made segmentation and deep learning pipelines for the classification of septin cages were written in ImageJ and Python using Keras.

### Statistics

All graphs were plotted and statistical analysis were performed using Prism. n.s: non-significant, *: p-value < 0.05, **: p-value < 0.01, ***: p-value < 0.001).

## Code availability

Custom ImageJ macros, phyton scripts and Jupyter notebooks were annotated and deposited in Github (https://github.com/ATLopezJimenez/Toolset-high-content-analysis-of-Shigella-infection).

## Acknowledgements

We thank members of the Mostowy and Brian Ho labs for helpful discussion. We thank François-Xavier Campbell-Valois for providing TSAR plasmids. We thank the instructors of the EMBL course Deep Learning for Image Analysis, in particular Prof. Pejman Rasti. A.T.L.J. was funded by the Swiss National Science Foundation Early Postdoc.Mobility Fellowship (P2GEP3_188277) and the European Union’s Horizon 2020 research and innovation program under the Marie Skłodowska - Curie grant agreement no. H2020-MSCA-IF-2020-895330. D.B. was supported by the Deutsche Forschungsgemeinschaft (DFG) Walter Benjamin Programme (BR 6637/1-1). G.Ö.G. was funded by the Human Frontier Science Program (LT000436/2021-L). This research in S. Mostowy laboratory was supported by a Wellcome Trust Senior Research Fellowship (206444/Z/17/Z) and European Research Council Consolidator Grant (772853 - ENTRAPMENT).

---

**Supplementary table S1.**
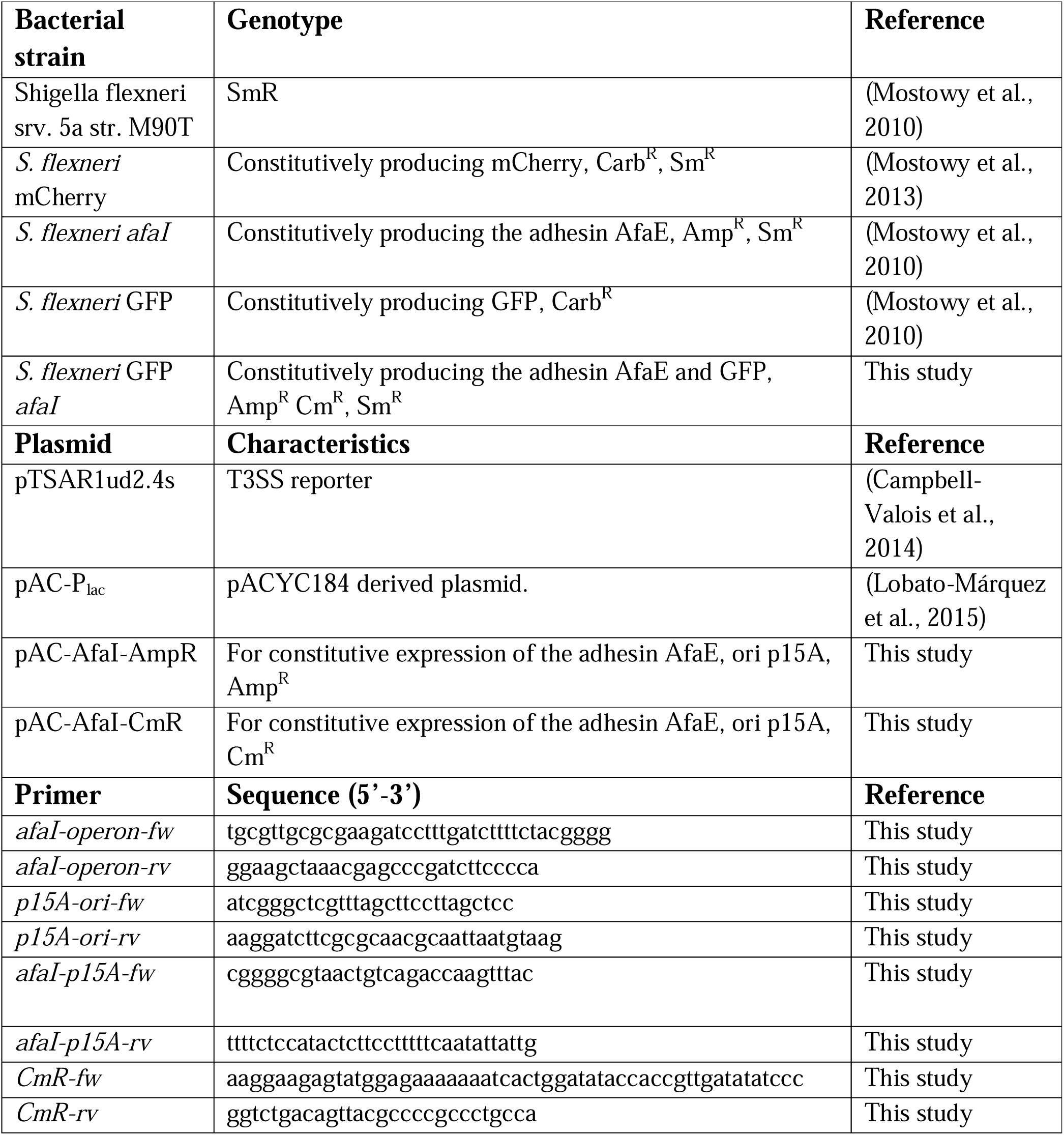

