## Supplementary Figures for "High-content high-resolution microscopy and deep learning assisted analysis reveals host and bacterial heterogeneity during *Shigella* infection"

**Figure 1-figure supplement 1**

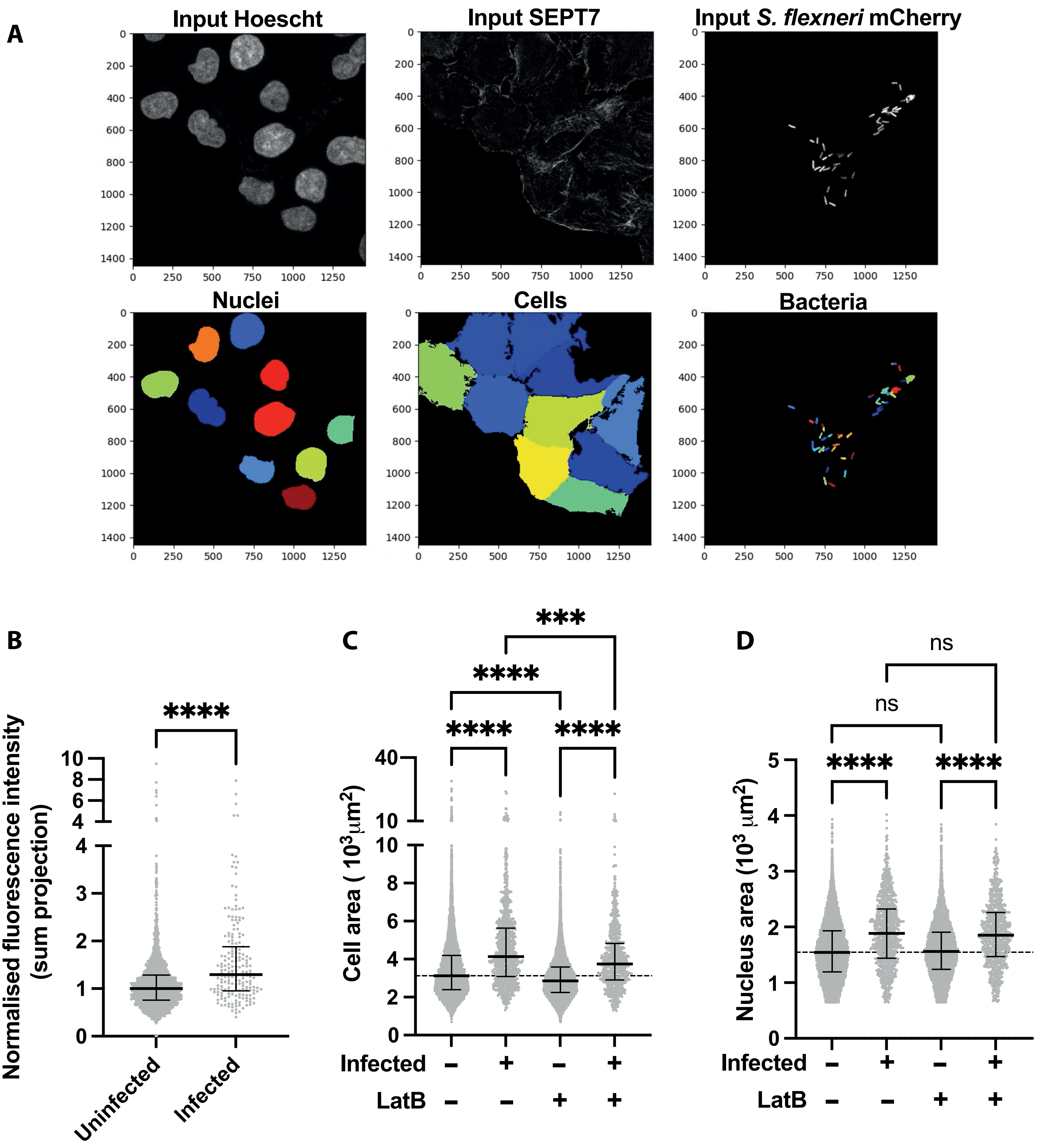

**Figure 2-figure supplement 1**

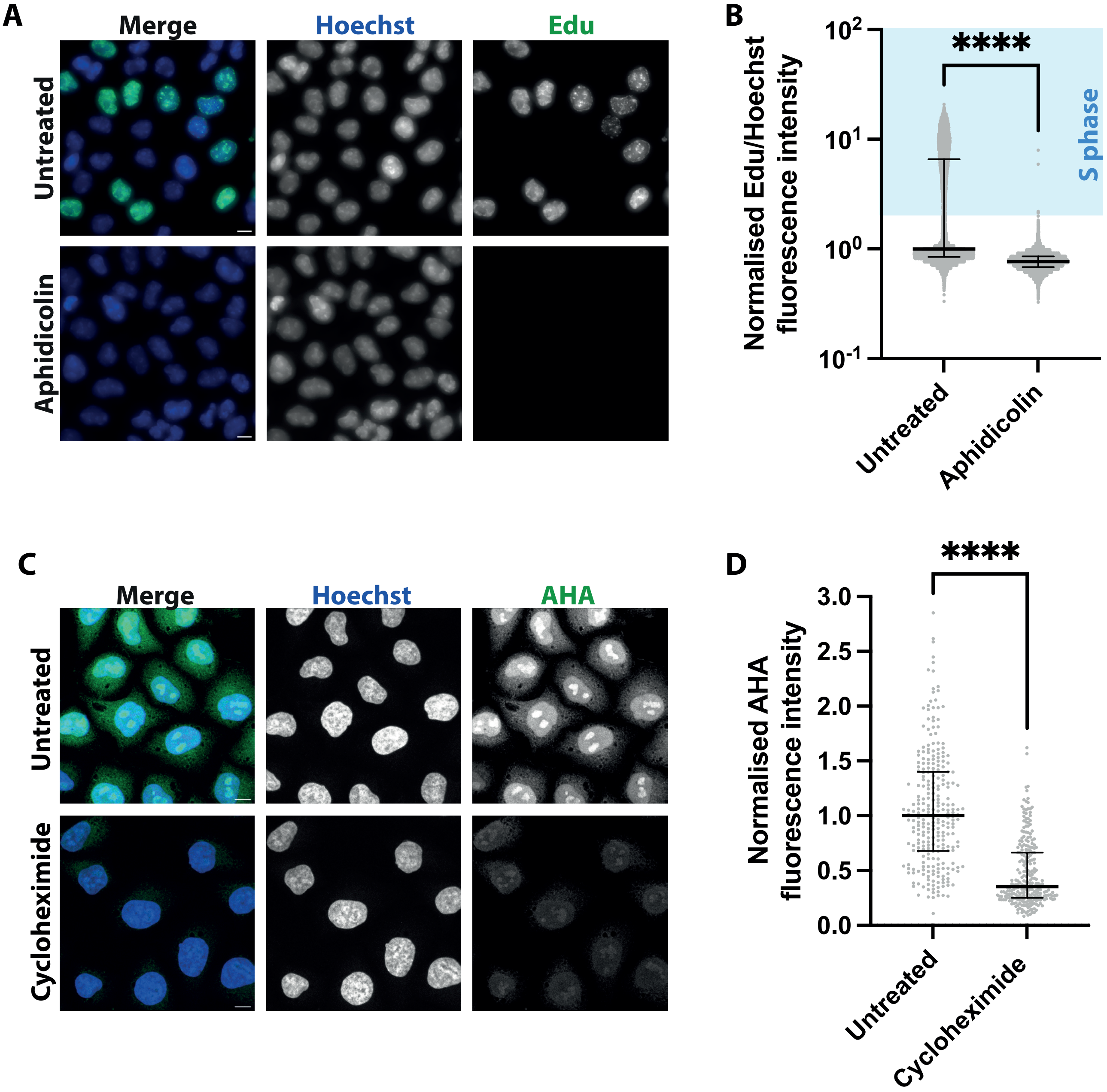

### Figure 4-figure suplement 1

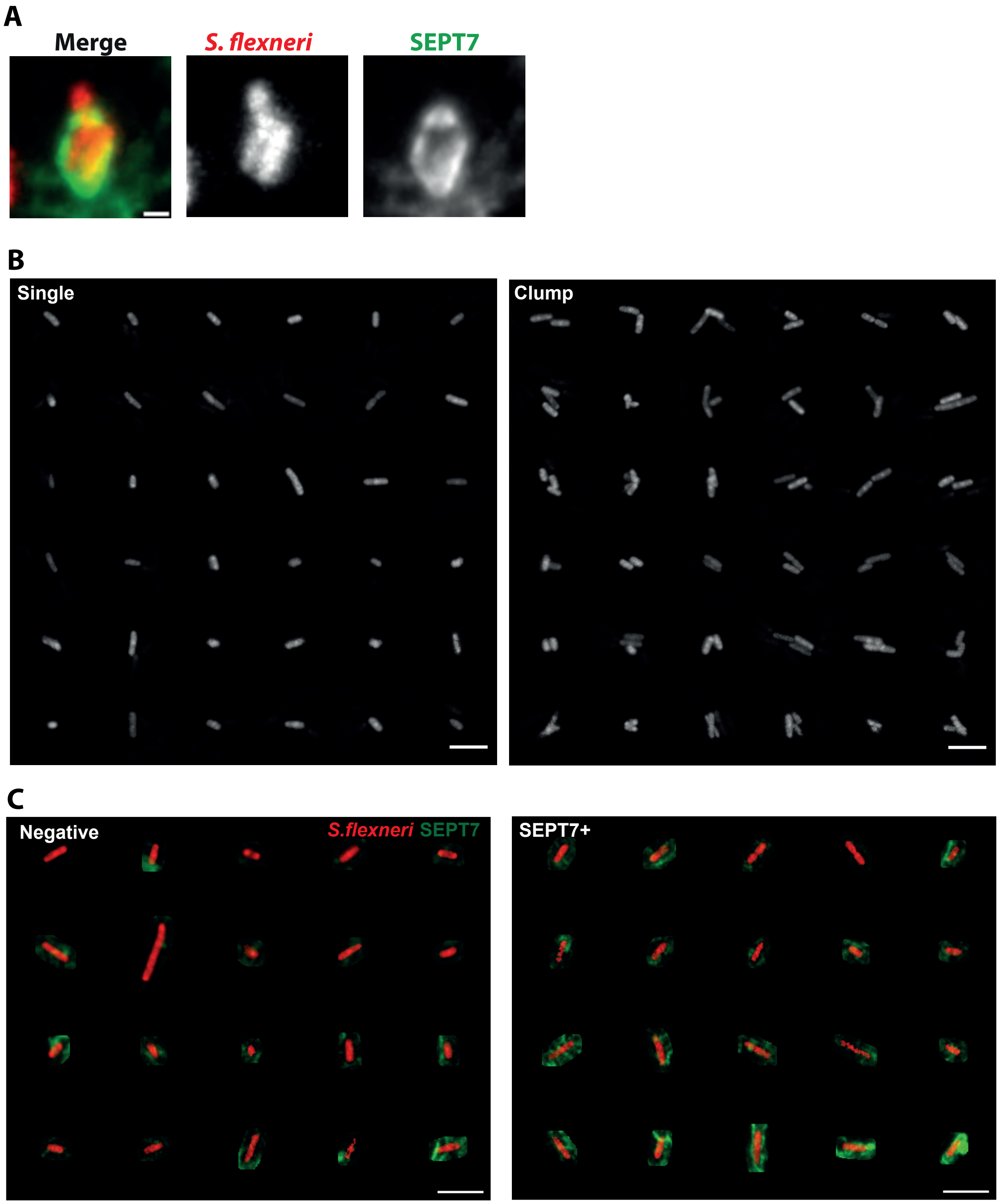

### Figure 5-figure supplement 1

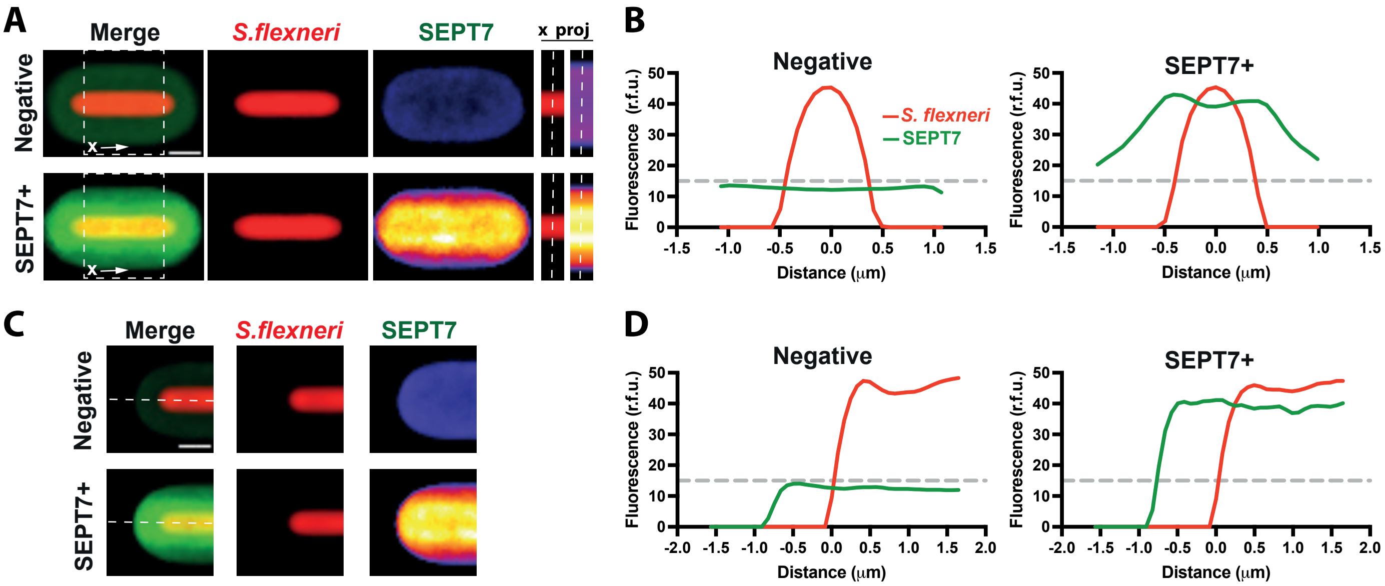

**Figure 6-figure supplement 1**

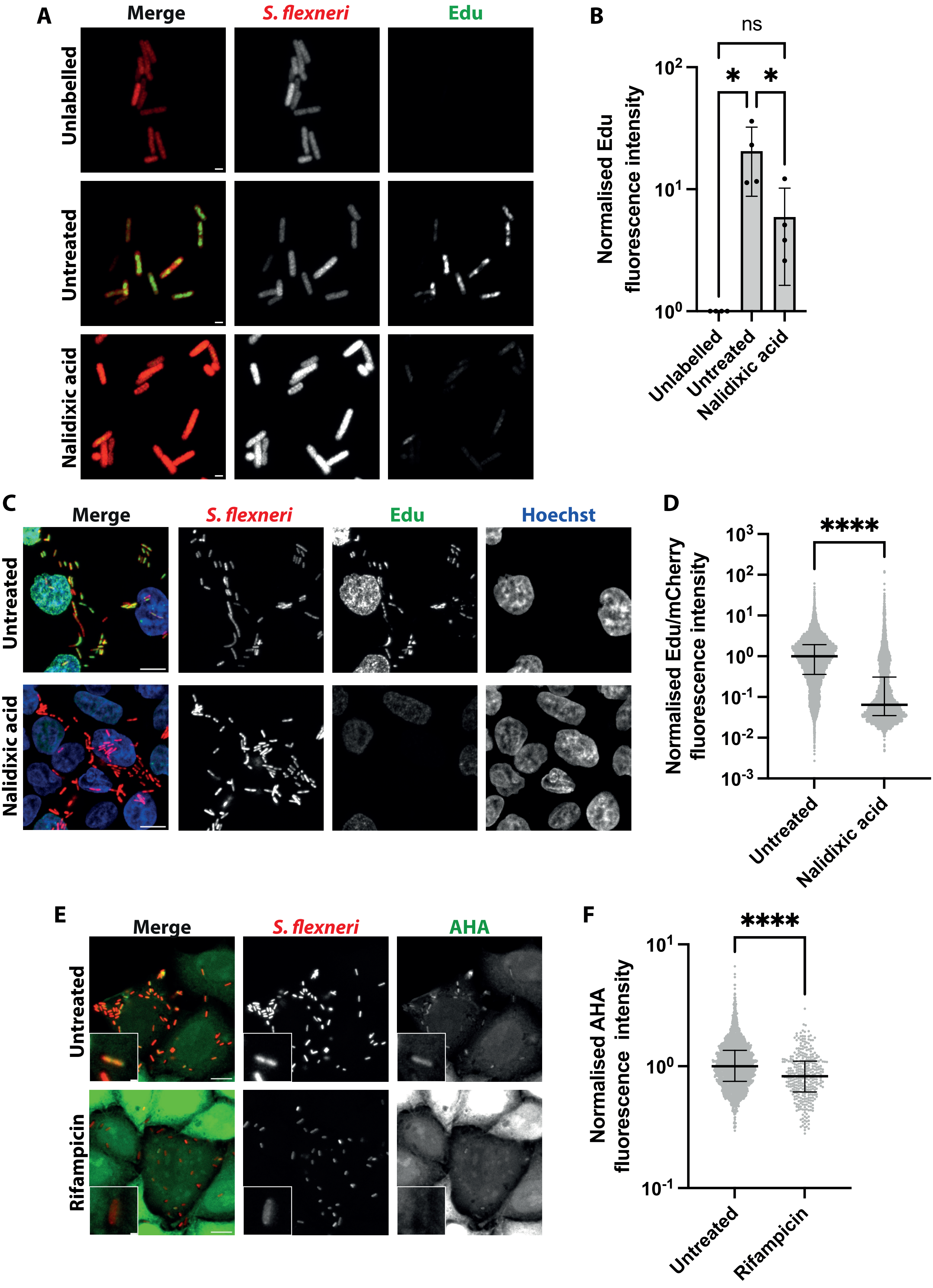

**Figure 7-figure supplement 1**

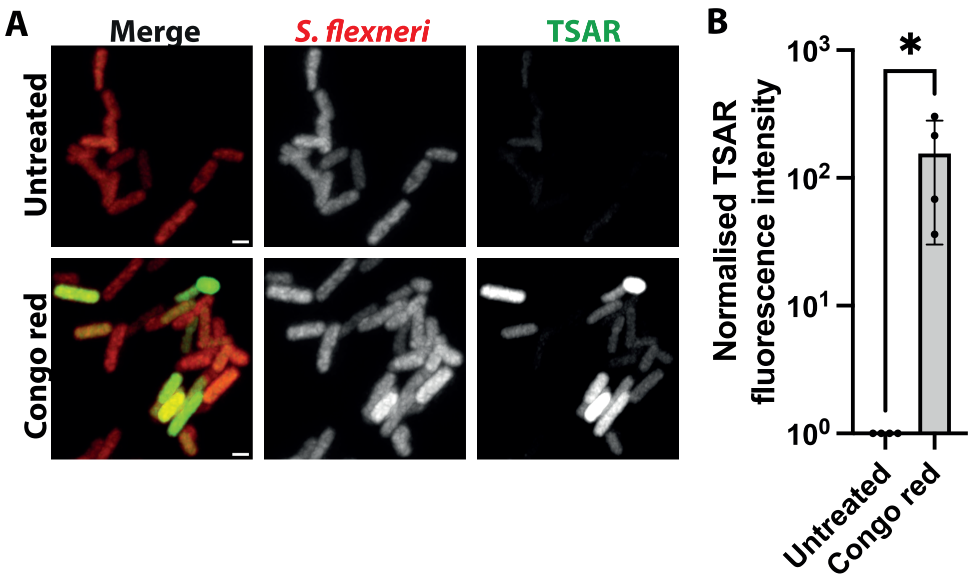


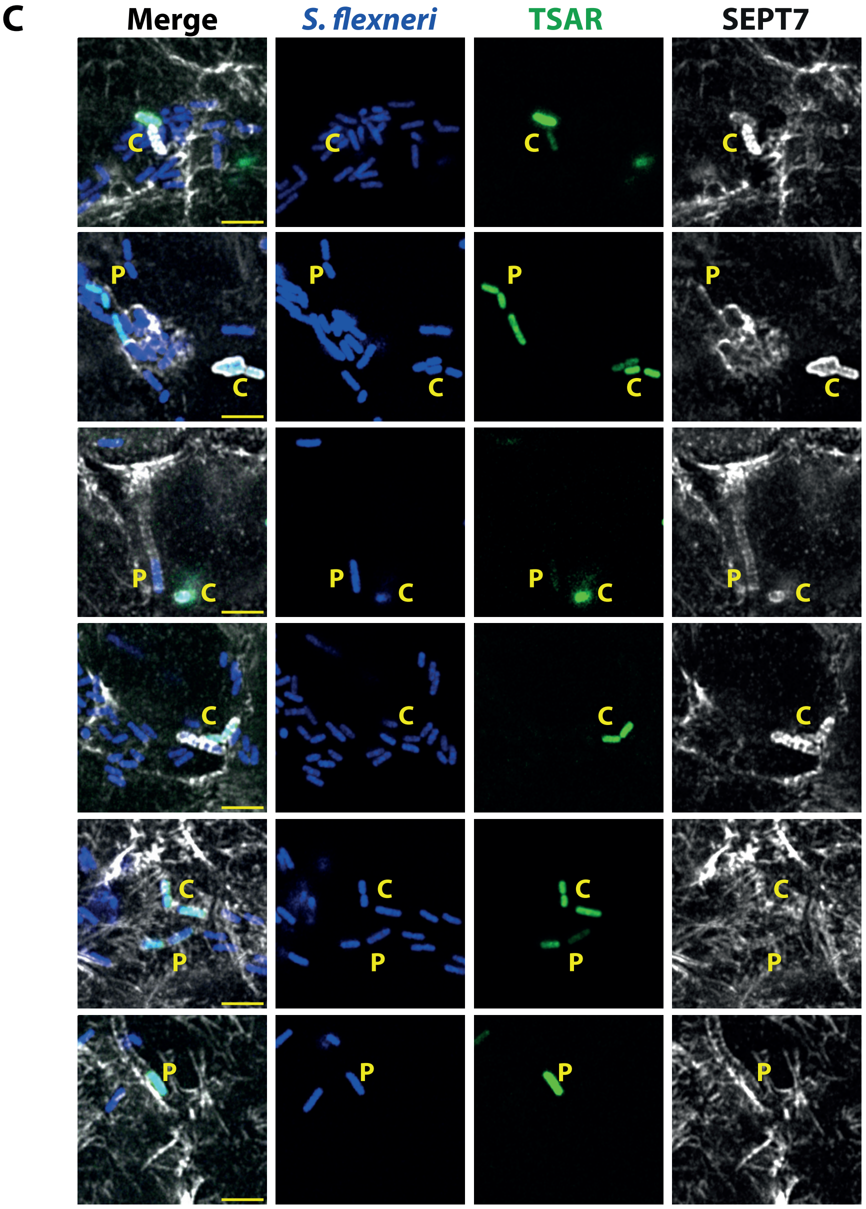
